# Genome-wide ChIPseq analysis of AhR, COUP-TF, and HNF4 enrichment in TCDD-treated mouse liver

**DOI:** 10.1101/2021.06.18.448955

**Authors:** Giovan N. Cholico, Rance Nault, Timothy R. Zacharewski

## Abstract

The aryl hydrocarbon receptor (AhR) is a ligand-activated transcription factor known for mediating the toxicity of 2,3,7,8-tetrachlorodibenzo-*p*-dioxin (TCDD) and related compounds. Although the canonical mechanism of AhR activation involves heterodimerization with the aryl hydrocarbon receptor nuclear translocator, other transcriptional regulators that interact with AhR have been identified. Enrichment analysis of motifs in AhR-bound genomic regions implicated co-operation with COUP transcription factor (COUP-TF) and hepatocyte nuclear factor 4 (HNF4). The present study investigated AhR, HNF4α and COUP-TFII genomic binding and effects on gene expression associated with liver-specific function and cell differentiation in response to TCDD. Hepatic ChIPseq data from male C57BL/6 mice at 2 hrs after oral gavage with 30 µg/kg TCDD were integrated with bulk RNA-sequencing (RNAseq) time-course (2 - 72 hrs) and dose-response (0.01 - 30 µg/kg) datasets to assess putative AhR, HNF4α and COUP-TFII interactions associated with differential gene expression. TCDD treatment resulted in the genomic enrichment of 23,701, 11,688, and 9,547 binding regions for AhR, COUP-TFII and HNF4α, respectively, throughout the genome. Functional enrichment analysis of differentially expressed genes (DEGs) identified differential binding enrichment for AhR, COUP-TFII, and HNF4a to regions within liver-specific genes suggesting intersections associated with the loss of liver-specific functions and hepatocyte differentiation. Analysis found that the repression of liver-specific, HNF4α target and hepatocyte differentiation genes, involved increased AhR and HNF4α binding with decreased COUP-TFII binding. Collectively, these results suggested TCDD-elicited loss of liver-specific functions and markers of hepatocyte differentiation involved interactions between AhR, COUP-TFII and HNF4α.

## INTRODUCTION

The aryl hydrocarbon receptor (AhR) is a ligand-activated transcription factor that belongs to the Per-Arnt-Sim basic-helix-loop-helix (PAS-bHLH) superfamily of proteins. When inactive, cytosolic AhR resides in the cytosol interacting with HSP90, AIP, p23 and SRC chaperone proteins (Rothhammer and Quintana, 2019). Following ligand binding, the chaperone proteins are shed and the AhR translocates into the nucleus. Canonical activation involves heterodimerization with the AhR nuclear translocator protein (ARNT) with subsequent binding to dioxin response elements (DREs) to elicit gene transcription (Rothhammer and Quintana, 2019). However, not all differentially expressed genes (DEGs) possess DREs, and studies have shown the AhR can bind to *non-consensus* DNA motifs to regulate gene expression (Denison et al., 2011; Dere et al., 2011). Moreover, the AhR can also interact with other transcription factors including the estrogen receptor (ER-α/β) (Ohtake et al., 2003), retinoic acid receptor (RAR) (Murphy et al., 2004), and Krüppel-like factor 6 (KLF6) (Wilson et al., 2013).

Although its endogenous function has yet to be resolved, numerous structurally diverse endogenous metabolites, natural products, and microbial compounds bind and activate the AhR (Larigot et al., 2018). The AhR also mediates effects in response to polycyclic aromatic hydrocarbons (PAHs), polychlorinated dibenzo-*p-* dioxins (PCDDs), dibenzofurans (PCDFs), and biphenyls (PCBs) (Larigot et al., 2018). 2,3,7,8-Tetrachlorodibenzo-*p*-dioxin (TCDD) is the most potent AhR ligand known to elicit a plethora of adverse effects across a broad spectrum of tissues and species. For example, an acute oral dose of TCDD induces hepatic steatosis with extrahepatic immune cell infiltration (Boverhof et al., 2005), with chronic exposure resulting in the progression of fatty liver to steatohepatitis with fibrosis (Pierre et al., 2014). Although the mechanism of TCDD and related compound toxicity is poorly understood, knockout studies indicate that most, if not all, adverse effects are the result of differential gene expression mediated by the AhR (Fernandez-Salguero et al., 1996).

The AhR has also been implicated in indirectly cooperating with other transcriptional regulators. AhR-bound DNA regions following TCDD treatment in the mouse liver are enriched in binding motifs for hepatocyte nuclear factor 4 (HNF4), COUP transcription factor (COUP-TF), peroxisome proliferator-activated receptor (PPAR), retinoid X receptor (RXR), liver X receptor (LXR), and nuclear factor erythroid 2-related factor 2 (NRF2) (Dere et al., 2011; Nault et al., 2018). These transcription factors regulate hallmark hepatic functions including lipid metabolism, antioxidant responses, and hepatocyte differentiation, all functions dysregulated by TCDD (Angrish et al., 2013, 2012; Cholico et al., 2021; Doskey et al., 2020; Fader et al., 2017b, 2015; Lu et al., 2011; Nault et al., 2018, 2017a, 2017b; Yeager et al., 2009). Moreover, treatment with TCDD every 4 days for 28 days resulted in a loss of liver-specific gene expression leaving a functionally dedifferentiated hepatocyte phenotype (Nault et al., 2017a).

Hepatocyte differentiation is primarily mediated by HNF4α as early as endoderm formation during embryogenesis (Parviz et al., 2003). HNF4α maintains differentiation by regulating hallmark hepatocyte functions such as lipid and bile metabolism, gluconeogenesis, amino acid metabolism, and blood coagulation (Bonzo et al., 2012). HNF4α is a nuclear transcription factor that regulates gene expression as a homodimer but can involve more complex interactions such as heterodimerization with COUP-TFs and RXR. Specifically, HNF4α co-interaction with COUP-TFII elicits tissue-specific inhibitory and synergistic gene expression effects (Ashraf et al., 2019). Alternatively, COUP-TFII dimerizes with RXR and competes with HNF4α/RXR for the same binding site or sequesters RXR thereby limiting HNF4α/RXR signaling events (Ashraf et al., 2019). Considering that COUP-TFs can functionally inhibit HNF4α-signaling in the liver, and that AhR-genomic binding motifs are enriched with HNF4α binding sites, we tested the hypothesis that AhR, HNF4α, and COUP-TFII potentially intersect to regulate liver-specific functions and hepatocyte differentiation following TCDD treatment.

## METHODS

### Animal Treatment

Male C57BL/6 mice (Charles River Laboratories, Kingston, NY) were treated as previously described (Nault et al., 2018). Briefly, mice were housed at 23°C, with 30-40% humidity, in 12 hr light/dark cycles. Food and water were available to mice *ad libitum*. TCDD (AccuStandard, New Haven, CT) was dissolved in acetone and diluted in sesame oil to a working stock. Postnatal day 28 mice were weighed and treated with 30 µg/kg TCDD or sesame oil. Mice were asphyxiated with carbon dioxide at 2 hrs post-treatment. The liver tissues were excised and snap frozen in liquid nitrogen. Tissues were then stored at −80°C until ChIP-sequencing (ChIPseq) could be conducted. Five mice were used for each treatment group. Animal studies were conducted with the approval of the Michigan State University Institutional Animal Care and Use Committee (PROTO201800043) and meet ARRIVE guidelines.

### ChIP-Sequencing

ChIPseq analysis of COUP-TFII and HNF4α was conducted through Active Motif (Carlsbad, CA). Equal amounts of liver tissue following a 2 hrs 30 μg/kg TCDD or vehicle exposure from five C57BL/6 male mice (∼100 mg) were pooled prior to ChIPseq analysis. Previously published ChIPseq data was used to identify AhR genomic binding sites after 2 hrs of 30 μg/kg TCDD exposure in C57BL/6 male mice (Fader et al., 2017b)(GEO: GSE97634). The fold-change for each genomic binding site was determined by comparing peak values from TCDD-treated samples against the vehicle control samples. Peaks with a 0.6 ≥ fold-change ≥ 1.5 were considered differentially enriched. The ChIPseq data for COUP-TFII and HNF4α have been deposited on the Gene Expression Omnibus (GEO; GSE178168).

### RNA-Sequencing

Previously published time-dependent (Nault et al., 2018) (GEO: GSE109863) and repeated-dose dose-response (Fader et al., 2017a)(GEO: GSE87519), hepatic RNAseq datasets were used to assess transcriptional changes following TCDD exposure in mice. Briefly, for the time-dependent study, mice were exposed to 30 μg/kg TCDD (or vehicle control) for 2, 4, 8, 12, 24, or 72 hrs, after which liver tissue was collected and bulk RNAseq was performed. For the repeated-dose dose-response study, mice were exposed to 0.01, 0.03, 0.1, 0.3, 1, 3, 10, or 30 μg/kg TCDD (or vehicle control) ever 4 days for 28 days, after which liver tissue was collected and bulk RNAseq was performed. For both datasets, gene expression fold-changes were calculated between TCDD-treated and time-matched vehicle-treated mice, along with posterior probabilities [P1(*t*)]. Genes were considered differentially expressed with a 0.6 ≥ fold-change ≥ 1.5 and P1(*t*) ≥ 0.8.

### Functional and Motif Enrichment Analyses

Functional enrichment was performed using BC3NET (de Matos Simoes and Emmert-Streib, 2012). Gene sets obtained from the gene set knowledgebase (GSKB; http://ge-lab.org/gskb/) were filtered to include only terms from the following databases: SMPDB, GO, KEGG, REACTOME, EHMN, MICROCOSM, MIRTARBASE, MPO, PID, PANTHER, BIOCARTA, INOH, NETPATH, WIKIPATHWAYS, MOUSECYC, TF, TFACTS. The subset GSKB gene sets were then complemented with in-house derived gene sets. All gene sets that were assessed for functional enrichment can be found at that Harvard Dataverse (https://doi.org/10.7910/DVN/OCKYFO). Protein encoding genes defined in Ensembl GRCm38 (released 102) were used as the reference gene set (Durinck et al., 2009). Analysis of known DNA motifs was conducted for differentially enriched regions in each ChIPseq dataset using HOMER v4.11.1 (http://homer.ucsd.edu/homer/motif/). All enriched motifs can be found in the supplementary file.

### ChIPseq Transcription Factor Overrepresentation Analysis

Overrepresentation of AhR, COUP-TFII, and HNF4α binding patterns (e.g., increased, decreased, and no change) on subsets of differentially expressed genes (DEGs) was performed using BC3NET. Gene sets were determined by assessing all possible transcription factor binding patterns for intragenic regions within the AhR, COUP-TFII, and HNF4α ChIPseq data. All DEGs in the RNAseq data of mice treated with 30 μg/kg TCDD every 4 days for 28 days were used as the reference gene set.

### Gene Set Enrichment Analysis

GSEA (v 4.0.3) was used to perform a gene set enrichment analysis (GSEA) (Subramanian et al., 2005) using a pre-ranked list of genes based on fold-change (induction to repression) for mice treated with 30 μg/kg TCDD every 4 days for 28 days.

## RESULTS

### Evaluation of ChIPseq Datasets

ChIPseq analysis of COUP-TFII and HNF4α identified a total of 30,960 and 43,233 enriched genomic binding locations (**Fig. 1**). These data were analyzed in combination with published AhR ChIPseq data that reported 27,413 enriched genomic regions following treatment with 30 µg/kg TCDD for 2 hrs (Nault et al., 2016b). Binding was predominantly localized to intragenic (−10Kb of transcription start site to transcription stop site) regions for the AhR (57.4%), COUP-TFII (81.5%) and HNF4α (76.4%). Evaluation of TCDD-elicited changes in genomic binding showed only AhR binding increased (median Log_2_ fold-change = 0.99). Only positive Log_2_ fold-changes in genomic AhR enrichment were reported since the AhR is a ligand-activated transcription factor that predominantly resides in the cytosol. In contrast, COUP-TFII (median Log_2_ fold-change = −0.30) and HNF4α (median Log_2_ fold-change = 0.13) exhibited both increased and decreased binding following treatment with 30 µg/kg TCDD for 2 hrs.

**Figure 1:**
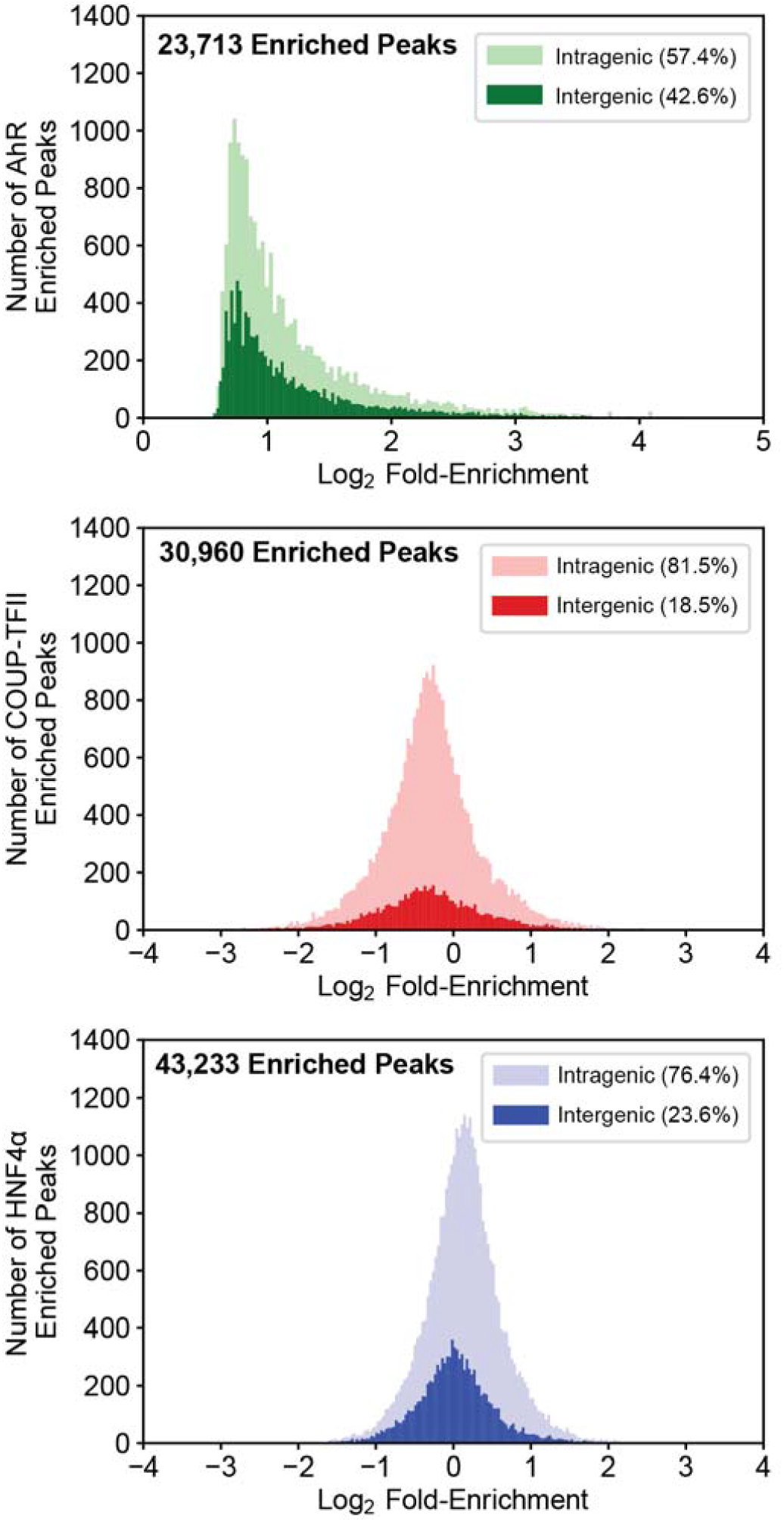
AhR, COUP-TFII, and HNF4α ChIPseq analysis in TCDD-treated liver samples. Mice were treated for 2 hrs with a single oral dose of 30 μg/kg TCDD (or sesame oil vehicle). Liver samples were assessed for AhR, COUP-TFII, and HNF4α genomic binding enrichment using ChIPseq. Log_2_ fold-enrichments were assessed for frequency of binding within intragenic (light color) and intergenic (dark color) regions. Intragenic regions are defined as 10Kb upstream of the transcriptional start site (TSS) to the end of the transcribed region of a gene. Intergenic is defined as regions outside of intragenic regions. The AhR histogram is depicted as a minimum log_2_ fold-enrichment of 0 as it does not constitutively bind DNA in the absence of a ligand.

Differential AhR, COUP-TFII and HNF4α ChIPseq peak enrichment, characterized by 0.6 ≥ fold-change ≥ 1.5, was examined (**Fig 2**.). Of the 23,713 AhR enriched peaks, 23,701 were defined as enriched (99.9%). In contrast, 11,688 of the 30,960 (37.8%) COUP-TFII peaks were enriched, and 9,547 of the 42,233 (22.6%) HNF4α peaks were enriched. COUP-TFII possessed 2,720 positively (i.e., increased binding) differentially enriched peaks and 8,968 negatively (i.e., reduced binding) differentially enriched peaks. HNF4α possessed 6,224 positively differentially enriched peaks and 3,323 negatively differentially enriched peaks. The sequences for these differentially enriched regions were assessed for over-represented DNA binding motifs. For AhR-enriched sequences, 31.53% and 13.06% possessed DNA binding motifs for AhR/Arnt and HNF4α, respectively, as well as COUP-TFII (25.10%; data not shown). Positively differentially enriched COUP-TFII sequences binding sites contained over-represented COUP-TFII (26.89% and 30.60%) and HNF4α (11.15%) binding motifs, while negatively differentially enriched COUP-TFII sequences possessed two different over-represented COUP-TFII binding sites (29.03% and 32.62%). Similarly, positively differentially enriched HNF4α sequences possessed over-represented HNF4α (7.42%), COUP-TFII (18.88% and 21.82%) and AhR/Arnt (10.71%) binding motifs, while negatively differentially enriched HNF4α sequences possessed over-represented HNF4α binding motifs (30.42%).

**Figure 2:**
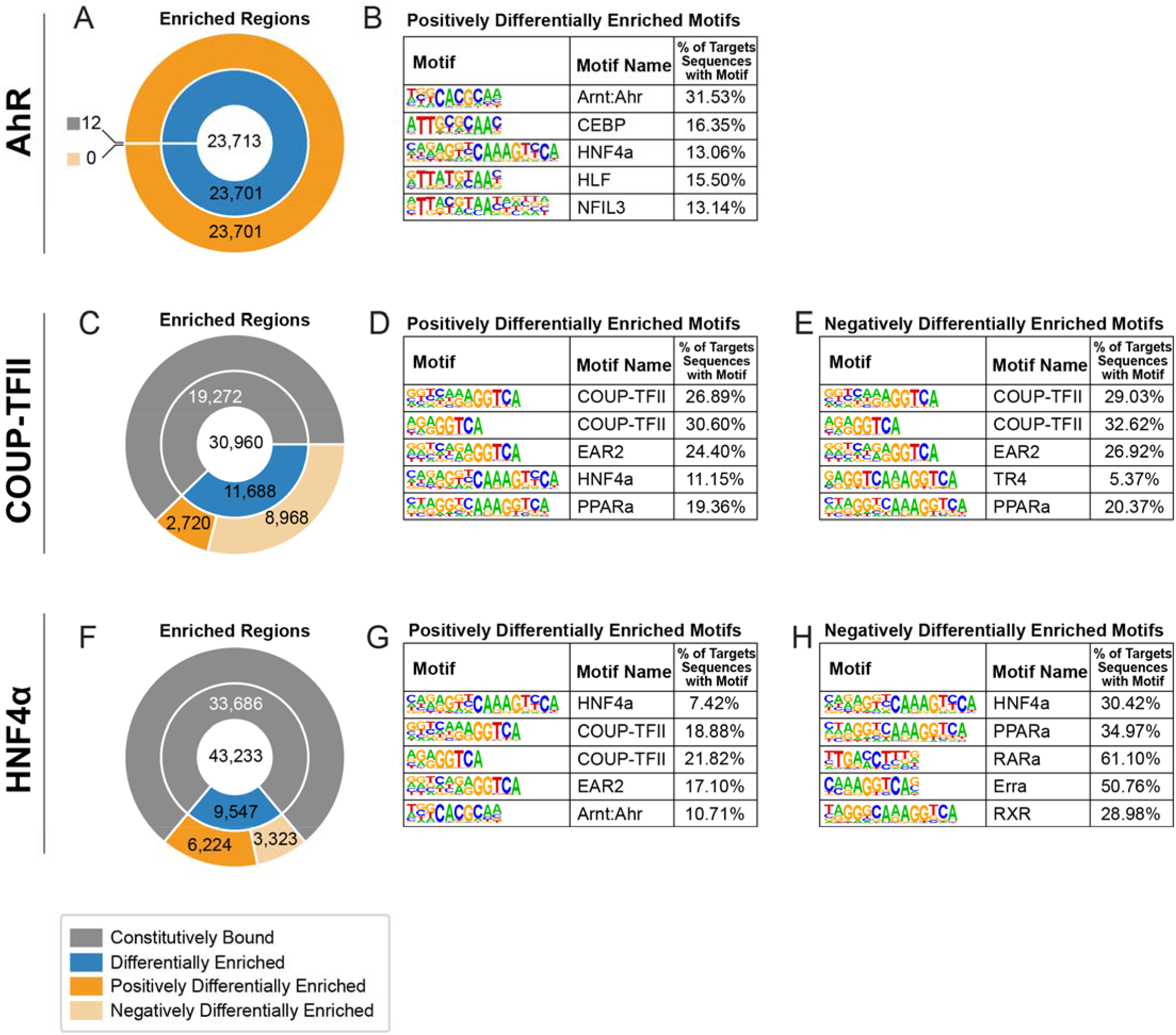
Over-represented motif analysis within differentially enriched peak sequences. **(A, C, F)** The effect of TCDD on the total number of enriched regions (center of donut plots) for AhR, COUP-TFII and HNF4a (0.6 ≥ fold-change ≥ 1.5; blue regions). Differentially enriched regions were further assessed for increased (positive – increased binding) and decreased (negative – decreased binding) enrichment (shades of orange). Sequences within positive **(B, D, G)** and negative **(E, H)** differentially enriched regions for each transcription factor were analyzed for enriched DNA motifs. The top 5 ranked binding motifs for each group are depicted. All over-represented motifs identified are listed in supplementary file 1.

Differentially enriched peaks between the three ChIPseq data sets were evaluated for overlap (**Fig. 3**). In reference to all differentially enriched AhR peaks, 47.85% and 70.09% of these peaks also overlapped with differentially enriched COUP-TFII and HNF4α peaks, respectively. In reference to differentially enriched COUP-TFII peaks, 25.57% and 72.87% overlapped with differentially enriched AhR and HNF4α peaks, respectively, while 28.15% and 53.93% of all differentially enriched HNF4α peaks overlapped with differentially enriched AhR and COUP-TFII peaks, respectively.

**Figure 3:**
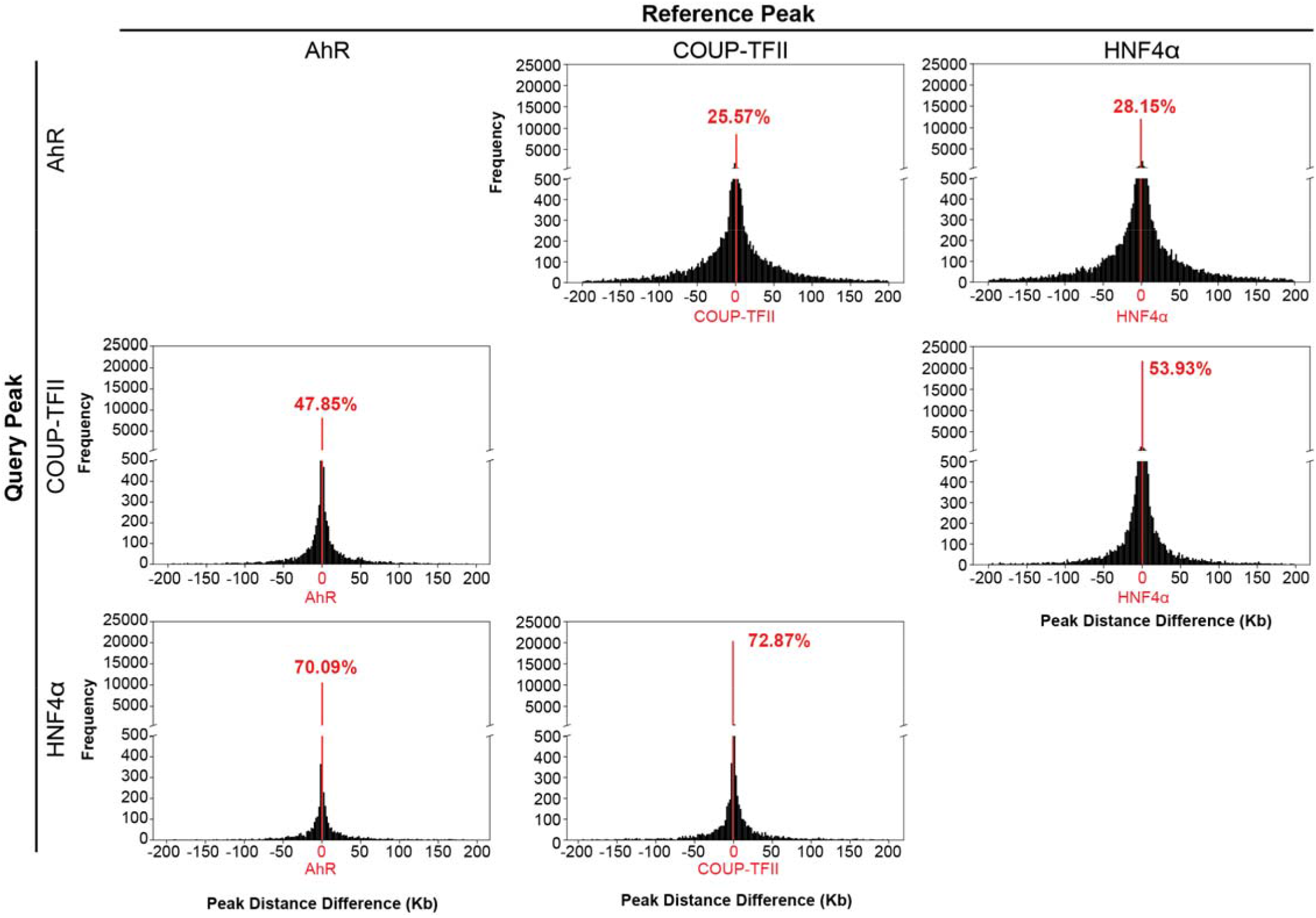
Evaluating overlap of differentially enriched peaks. Differentially enriched peaks for AhR, COUP-TFII and HNF4α were assessed for overlap following a 2 hr single oral gavage of 30 μg/kg TCDD. The distance (Kb) prevalence between transcription factor peak centers is denoted in each histogram. The number of overlapping peaks out of the total number of peaks for the reference transcription factor is denoted in red as a percentage.

### Evaluation of Transcription Factor Gene Targets

Intragenic peaks were further evaluated to identify putative gene targets (**Fig. 4**). Gene targets harboring binding sites for multiple transcription factors anywhere within the intragenic region (regardless of overlap between transcription factor binding sites) were defined as possibly possessing *intersections* between AhR, COUP-TFII and HNF4α in the present study. Gene targets were assessed for binding of individual transcription factors, the intersection between two of the transcription factors, and the intersection between all three transcription factors. 7,762, 13,377 and 13,744 unique genes were identified as possessing intragenic binding of AhR, COUP-TFII, and HNF4α, respectively. Putative effects of TCDD were determined by associating differentially enriched peaks with the closest gene within the intragenic region, thereby limiting the number of potential targets for AhR, COUP-TFII, and HNF4α to 7,761, 6,846, and 5,762 unique genes, respectively. Potential intersections between transcription factors and subsequent putative gene co-regulation were determined by identifying common genes in which at least one of three transcription factors possessed a differentially enriched peak. It was assumed that a change in binding for only one transcription factor is necessary to elicit differential gene expression. The number of genes showing co-enrichment of AhR and COUP-TFII was 6,687, while 6,954 and 8,080 showed co-enrichment with at least one differentially enriched transcription factor for AhR and HNF4α, or COUP-TFII and HNF4α, respectively. A total of 6,376 genes exhibited co-enrichment with at least one differentially enriched transcription factor for AhR, COUP-TFII and HNF4α.

**Figure 4:**
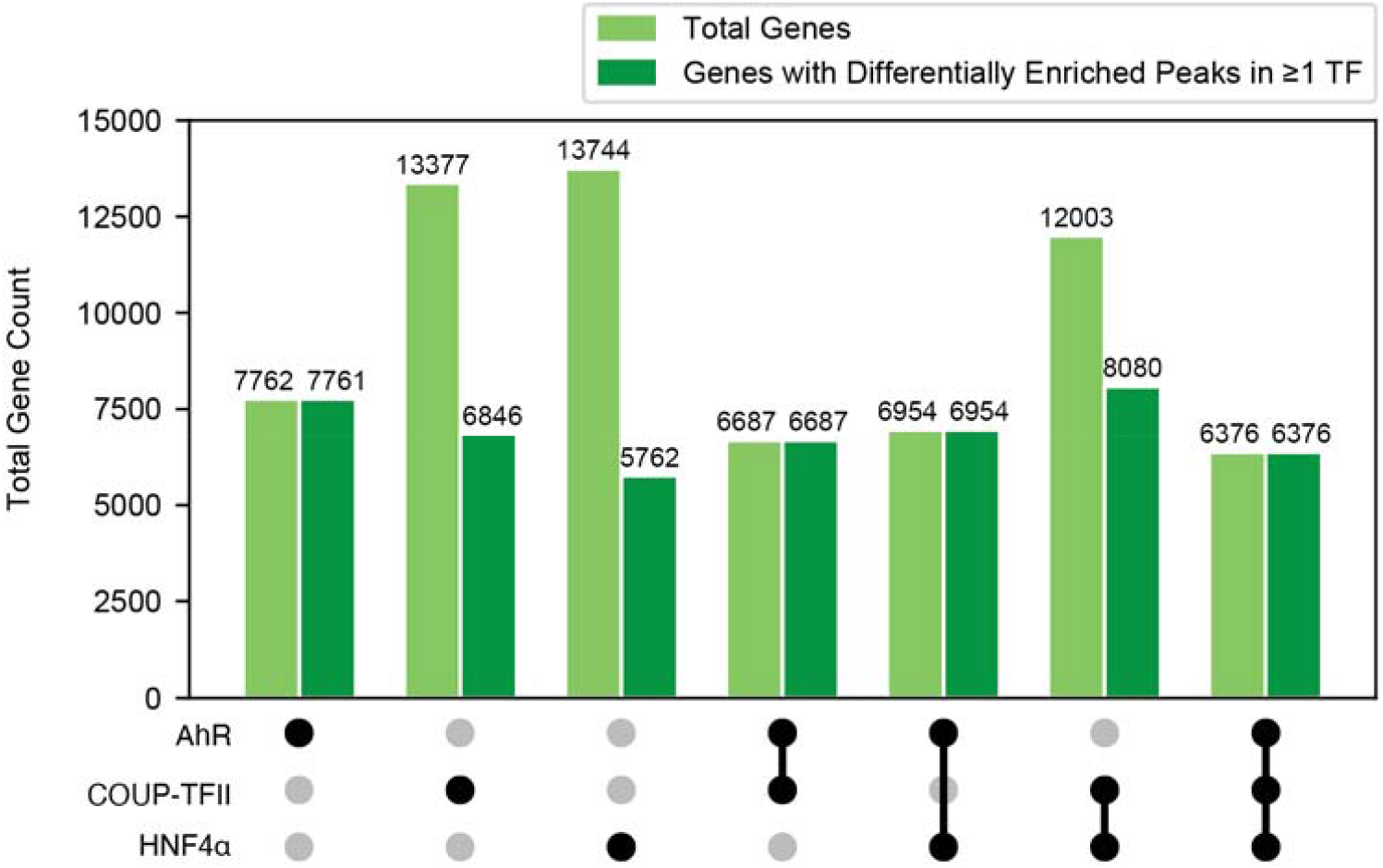
Evaluating putative co-regulated gene targets. The total number of hepatic genes associated with AhR, COUP-TFII and HNF4a enrichment in intragenic regions in ChIPseq data using mouse liver samples 2 hrs after TCDD treatment. Black circles denote the presence of genes for each transcription factor dataset. Common genes among transcription factor datasets are also identified and are referred to as “intersections”. The number of genes possessing at least one enriched peak following TCDD treatment are denoted in dark green.

### Evaluation of Gene Transcriptional Changes

ChIPseq and RNAseq data were integrated to identify changes in gene expression for putative co-regulated genes (**Fig. 5**). Time course RNAseq data, in which mice were treated with 30 µg/kg TCDD and sacrificed 2, 4, 8, 12, 24, and 72 hrs after treatment, were used to assess the acute effects of TCDD on hepatic gene expression. Dose-response RNAseq data were used to assess the sub-chronic hepatic effects of TCDD in mice treated with 0.01, 0.03, 0.1, 0.3, 1, 3, 10, or 30 µg/kg TCDD every 4 days for 28 days total. Among DEGs enriched for AhR, COUP-TFII, and/or HNF4α that exhibited differential expression [0.6 ≥ fold-change ≥ 1.5 and P1(*t*) ≥ 0.8] following acute exposure, the median fold-change was positive indicating primarily induction (**Fig. 5A**). Conversely, sub-chronic TCDD treatment elicited mixed differential expression in transcription factor-bound genes. For example, 0.01 to 3 µg/kg induced and repressed the expression of putatively co-regulated genes. In contrast, mice gavaged with 10 or 30 µg/kg exhibited more repression of putative co-regulated differentially expressed genes (DEGs). Of the 6,376 genes that were putatively co-regulated by AhR, COUP-TFII and HNF4α, 2,680 were differentially expressed (median Log_2_ fold-change = −0.79) after oral gavage every 4 days for 28 days with TCDD. Of these 2,680 DEGs, 1,106 were induced and 1,574 were repressed (**Fig. 5B**).

**Figure 5:**
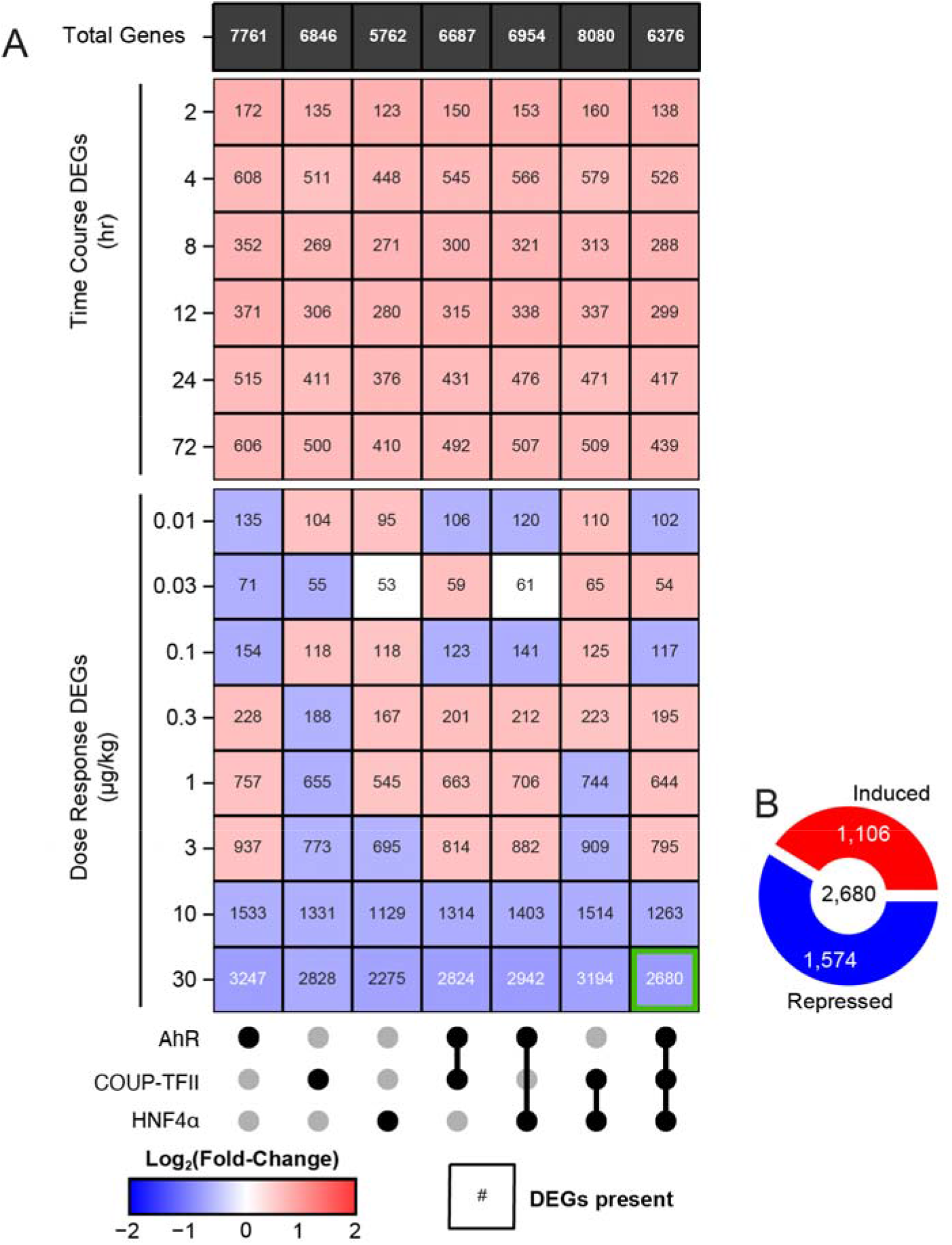
Gene expression of putative co-regulated genes. **(A)** The total number of genes with at least one enriched peak (black row) was determined for each transcription factor and their intersection (black dots). Differential gene expression (heatmap) was assessed for the transcription factor bound genes in a time course (GSE109863) and dose response (GSE87519) RNAseq dataset. The number within each tile provides the total number of differentially expressed genes (DEGs) for each individual transcription factor and their intersections. The color of the heatmap tile denotes the median Log_2_ fold-change of differentially expressed genes. **(B)** DEGs putatively co-regulated by AhR, COUP-TFII, and HNF4α in mice gavaged with 30 µg/kg every 4 days for 28 days were further evaluated (genes represented in green square). The number of induced and repressed DEGs are shown in red and blue, respectively.

### Functional Enrichment of Putatively AhR, COUP-TFII and HNF4α Co-Regulated Genes in Mice Treated with 30 µg/kg for 28 days

A total of 2,680 hepatic DEGs from mice treated with 30 µg/kg TCDD every 4 days for 28 days that exhibited AhR, COUP-TFII and HNF4α genomic binding (6,376) were assessed for functional enrichment (**Table 1; Supp. File 2**). Analysis indicated that effects on diurnal regulated genes were over-represented, with 1,222 diurnal DEGs exhibiting differential expression (**Table 1**), followed by markers for portal hepatocytes (326 DEGs), midcentral hepatocytes (292 DEGs) and central hepatocytes (203 DEGs).

**Table 1:** **Top 20 enriched terms of DEGs putatively co-regulated by AhR/COUP-TFII/HNF4α in mice treated with 30 μg/kg TCDD every 4 days for 28 days**

| TermID | DEGs | All Genes<br>in Term | Adj.<br>*p*-Value |
| --- | --- | --- | --- |
| TZ_DIURNAL_GENES | 1222 | 5613 | 0 |
| TZ_SCS_PORTALHEP | 326 | 581 | 6.91E-269 |
| TZ_SCS_MIDCENTRALHEP | 292 | 563 | 1.76E-226 |
| TZ_SCS_CENTRALHEP | 203 | 466 | 3.59E-136 |
| GO_CC_MM_MITOCHONDRION_Ensembl | 308 | 1328 | 6.80E-120 |
| KEGG_MM_METABOLIC_PATHWAYS_Ensembl | 290 | 1180 | 1.59E-119 |
| HEPG2_SECRETOME | 216 | 757 | 1.53E-101 |
| PHH_SECRETOME | 211 | 726 | 7.27E-101 |
| TZ_LIVER | 103 | 181 | 3.77E-83 |
| GO_CC_MM_CYTOSOL_Ensembl | 210 | 927 | 3.02E-78 |
| TZ_SCS_MIDPORTALHEP | 81 | 149 | 8.88E-63 |
| GO_MF_MM_ATP_BINDING_Ensembl | 240 | 1454 | 3.22E-61 |
| TZ_MALE_ENRICHED | 113 | 367 | 1.08E-55 |
| GO_BP_MM_METABOLIC_PROCESS_Ensembl | 131 | 521 | 2.05E-53 |
| GO_MF_MM_METAL_ION_BINDING_Ensembl | 168 | 871 | 1.64E-51 |
| TZ_SCS_MACROPHAGES | 171 | 904 | 2.96E-51 |
| GO_MF_MM_PROTEIN_HOMODIMERIZATION_ACTIVITY_Ensembl | 125 | 509 | 9.79E-50 |
| TZ_SCS_ENDOTHELIALCELLS | 119 | 461 | 9.79E-50 |
| GO_BP_MM_OXIDATION-REDUCTION_PROCESS_Ensembl | 128 | 535 | 1.25E-49 |

Furthermore, of the 726 genes associated with the primary human hepatocyte secretome, 211 were differentially expressed (81 repressed, 31 induced) with 103 DEGs (100 repressed, 3 induced) out of 181 being specific to the liver.

Putatively co-regulated liver-specific DEGs were examined further for AhR, COUP-TFII, and HNF4α intersections associated with the reported loss of the liver functional “phenotype” following TCDD treatment (Nault et al., 2017a). To assess the unique contributions of each transcription factor in modulating the expression of liver-specific genes, we further evaluated the 156 DEGs following treatment with 30 μg/kg TCDD every 4 days for 28 days (**Fig. 6A**). Analysis indicated AhR enrichment for 53.8% of liver-specific DEGs while COUPTFII and HNF4a binding within the intragenic region was more limited and included positive as well as negative differential enrichment when compared to a random list of DEGs in the RNAseq dataset. This suggests possible cooperation between AhR, COUP-TFII, and HNF4α in regulating differential gene expression following TCDD treatment (**Fig. 6B**). Of the ranked over-representation analysis, 62 DEGs possessed an increase of AhR binding, a decrease of COUP-TFII binding, and no change in HNF4α binding was the most over-represented binding pattern for liver-specific DEGs (*p*-value = 6.78E-09; **Fig. 6B**). In addition, increased AhR and HNF4α binding, with and without decreased COUP-TFII (48 and 33 DEGs, respectively) were also a highly ranked binding patterns within liver-specific DEGs. ChIPseq data were mapped onto RNAseq data for the 156 liver-specific DEGs (**Fig. 6C**). Of the 156 liver-specific DEGs, only four (*Cyp1a2, Igfbp1, Ephx1, Ugt2b35*) were induced following treatment with 30 µg/kg TCDD. Integrated ChIPseq and RNAseq data suggests that dose-dependent repression of almost all liver-specific DEGs correlates with an increase in AhR and HNF4α binding, and a decrease in COUP-TFII binding. The effect of TCDD on the hepatocyte secretome gene set was also examined of loss of liver differentiation, which includes assessing genes associated with liver function. This includes assessing the genes encoding the liver secretome to evaluate hepatocyte changes in response to TCDD, as well as assess HNF4α-dependent genes since liver differentiation relies so heavily on HNF4α signaling.

**Figure 6:**
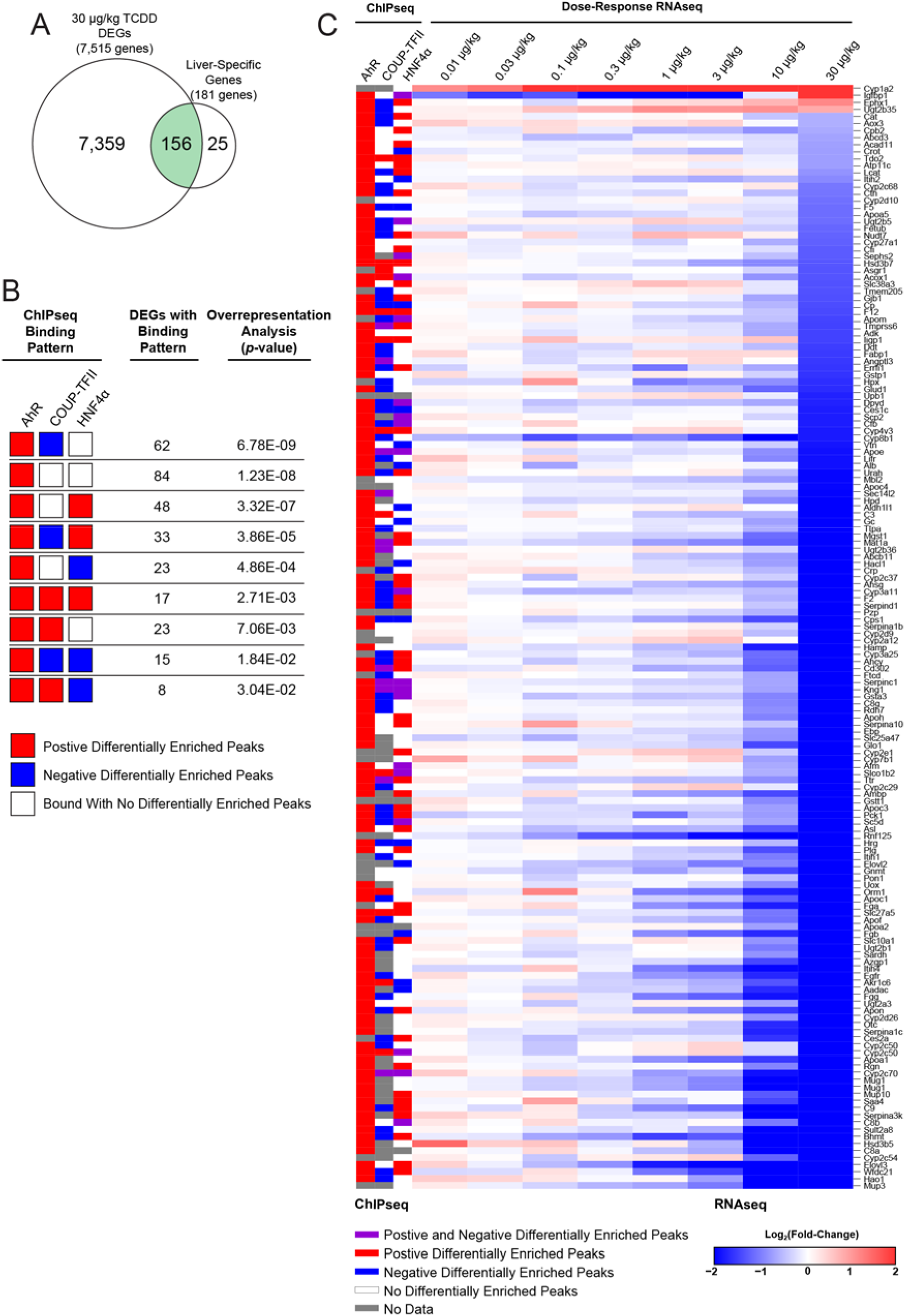
Integration of ChIPseq and dose response RNAseq for liver-specific DEGs. A total of 181 liver-specific genes were identified based on multi-tissue data from BioGPS as reported in Nault et al (Nault *et al*., 2017). **(A)** Differential gene expression was assessed for liver-specific genes in mice treated with 30 μg/kg TCDD every 4 days for 28 days. **(B)** Overrepresentation of transcription factor binding patterns for the 156 liver-specific DEGs was assessed using a one-sided Fisher’s exact test with FDR correction. **(C)** AhR, COUP-TFII, and HNF4α binding was mapped to expression of 103 liver-specific DEGs at all doses (0.01 - 30 µg/kg). Genes are ranked from most induced to most repressed at 30 μg/kg.

### Identification of AhR, COUP-TFII and HNF4α Binding Patterns in the Liver Secretome

Hepatocytes are responsible for the production and secretion of diverse proteins such as hepatokines, plasma proteins, and coagulation factors. Previous studies have identified 691 potentially secreted proteins in primary human hepatocytes (PHHs) using liquid-chromatography tandem mass spectrometry (Franko et al., 2019). Human genes encoding these 691 proteins were mapped to 726 mouse orthologues using biomaRt (Durinck et al., 2009). GSEA of mouse orthologues revealed that the PHH gene set was largely repressed (normalized enrichment score = −2.32) by 30 μg/kg TCDD every 4 days for 28 days (**Fig. 7A**). Of the 726 mouse PHH orthologues, 383 were differentially expressed by TCDD (**Fig 7B**). The binding pattern for PHH secretome DEGs favored an increase in AhR and HNF4α binding, and a decrease in COUP-TFII binding following TCDD treatment (**Fig 7C**). Specifically, 180 DEGs only exhibited an increase in AhR binding (*p*-value = 5.92E-11), followed by the DEGs with increased AhR binding and decreased COUP-TFII binding (113 genes; *p*-value = 5.43E-07) or increased HNF4α binding (79 genes; *p*-value = 8.45E-04). Collectively, the integration of ChIPseq and RNAseq data suggested that TCDD-elicited PHH secretome gene repression trended toward an increase in AhR and HNF4α binding and a decrease of COUP-TFII binding events.

**Figure 7:**
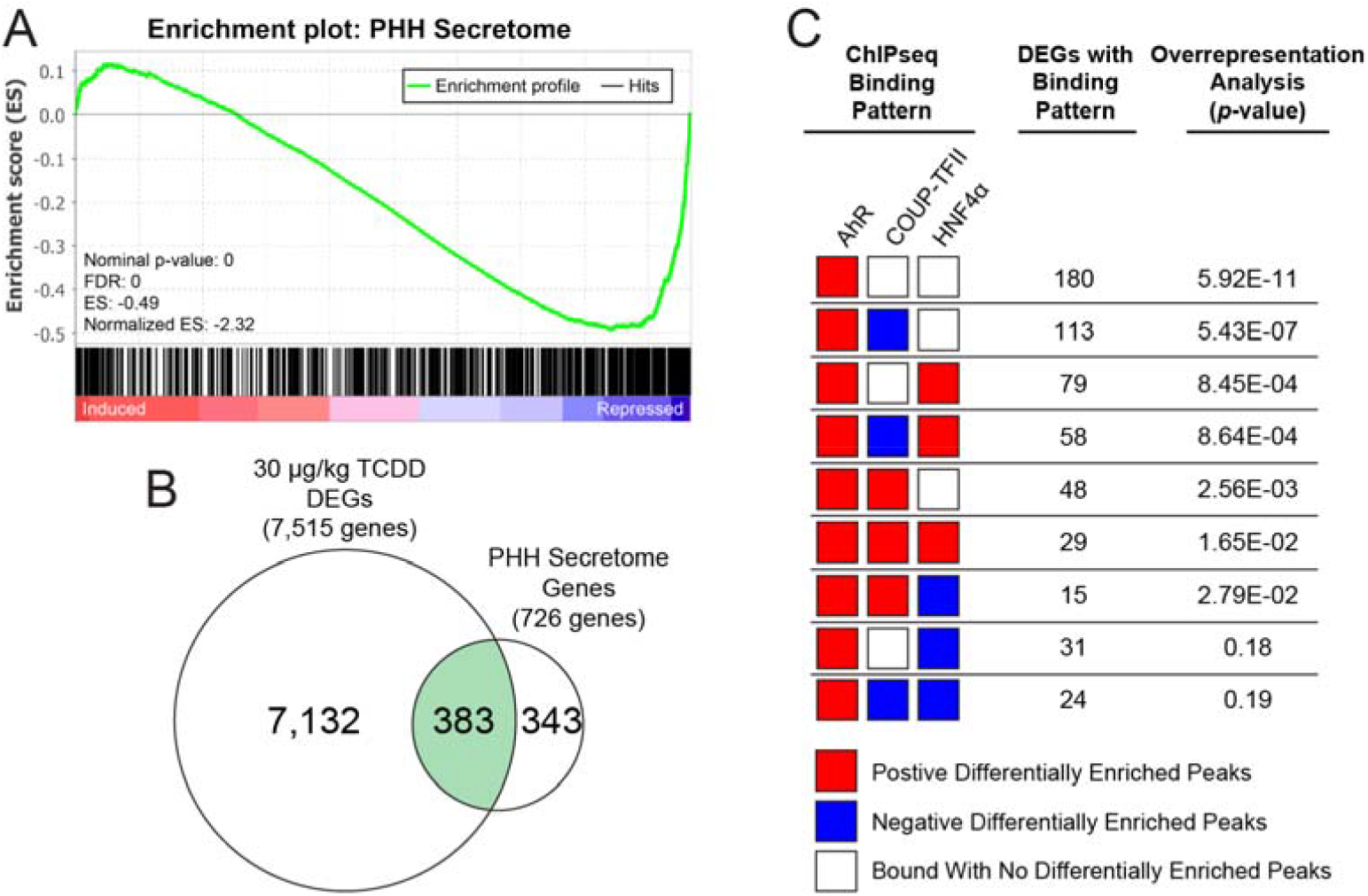
TCDD dose-dependently repressed expression of liver-specific secretome genes. Loss of liver function was assessed by examining changes in hepatic secretome gene expression. A putative primary human hepatocyte (PHH) secretome of 691 genes has been previously reported that was mapped to 726 mouse orthologues (Franko *et al*., 2019). **(A)** A gene set enrichment analysis for 726 mouse orthologs of the human PHH secretome was conducted using data for C57Bl/6 male mice orally gavaged with 30 ug/kg TCDD every 4 days for 28 days. **(B)** Differential gene expression was assessed for genes encoding the PHH secretome in mice treated with 30 μg/kg TCDD every 4 days for 28 days. **(C)** Over-representation of transcription factor binding patterns for the 383 PHH secretome DEGs was assessed using a one-sided Fisher’s exact test with FDR correction.

### Identification of AhR, COUP-TFII and HNF4α Binding Patterns in HNF4α-Dependent Genes

HNF4α is a key regulator of liver development (Parviz et al., 2003) and plays a pivotal role in liver homeostasis (Bonzo et al., 2012). Hepatocyte-specific ablation of HNF4α in mice resulted in the identification of 877 DEGs confirming the importance of hepatic HNF4α-signaling (Walesky et al., 2013). To assess the effect of TCDD on hepatocyte differentiation and function, GSEA of 576 hepatocyte-specific HNF4α-dependent genes revealed an overall repression in mice treated with 30 μg/kg TCDD every 4 days for 28 days (normalized enrichment score = −2.64; **Fig. 8A, B**). Over-representation of transcription factor binding patterns for these 576 genes identified an increase in AhR binding (252 genes; *p*-value = 4.30E-11), followed by increased AhR and decreased COUP-TFII binding (166 genes; *p*-value = 5.43E-07), or increased AhR and increased COUP-TFII binding (75 genes; *p*-value = 8.45E-04) (**Fig. 8C**). 109 DEGs exhibited an increase in AhR and COUP-TFII binding (*p*-value = 1.36E-03). Collectively, the repression of HNF4α-dependent genes by TCDD trended toward an increase in AhR with mixed effects on COUP-TFII binding events.

**Figure 8:**
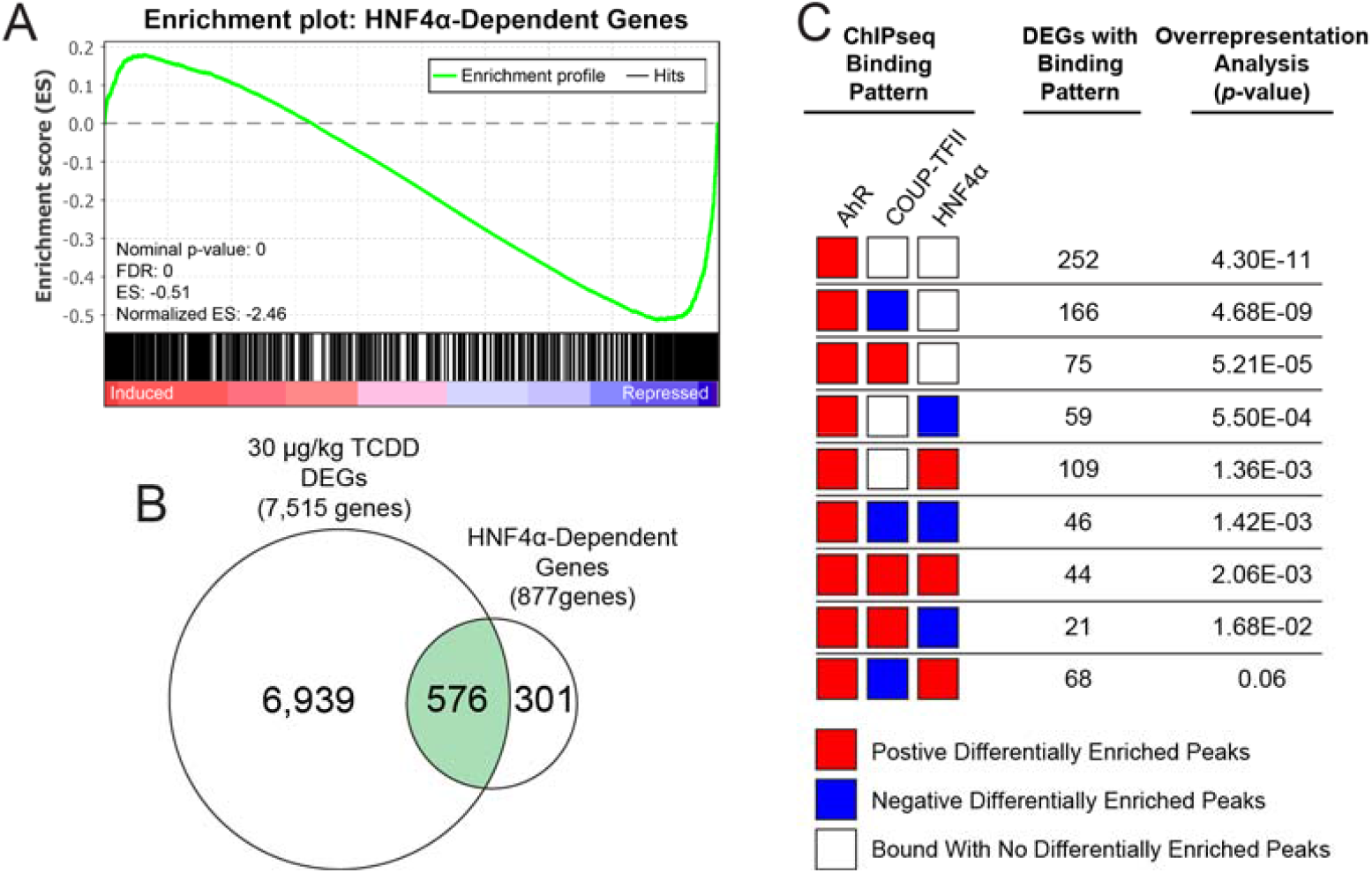
TCDD dose-dependently represses expression of HNF4α-dependent gene expression in the liver. Loss-of-liver function was assessed by examining changes in HNF4α-dependent gene expression. Putative hepatic HNF4α-dependent genes were previously identified (Walesky *et al*., 2013). **(A)** Gene set enrichment analysis for HNF4α-dependent genes was conducted using data for C57Bl/6 male mice orally gavaged with 30 ug/kg TCDD every 4 days for 28 days. **(B)** Differential expression was assessed for HNF4α-dependent genes in mice treated with 30 μg/kg TCDD every 4 days for 28 days. **(C)** Over-representation of transcription factor binding patterns for the 576 hepatic HNF4α-dependent DEGs was assessed using a one-sided Fisher’s exact test with FDR correction.

## DISCUSSION

The present study examined the effect of TCDD on AhR, HNF4α and COUP-TFII genomic binding with subsequent effects on the expression of genes associated with liver function and hepatocyte differentiation. As a potent AhR agonist, TCDD dysregulates a plethora of hepatic functions in rodents including lipoprotein assembly and export metabolism (Angrish et al., 2013; Nault et al., 2017b), bile acid homeostasis (Fader et al., 2017b; Forgacs et al., 2012), cholesterol metabolism (Angrish et al., 2013; Nault et al., 2017b), lipid metabolism (Angrish et al., 2013; Cholico et al., 2021; Nault et al., 2017b), glucose metabolism (Fader et al., 2019; Nault et al., 2016a, 2016b), iron and heme homeostasis (Fader et al., 2017a), one-carbon metabolism (Fling et al., 2020), and cobalamin-dependent reactions (Orlowska et al., 2021). Previous studies have established that the activated AhR can bind to DNA motif as a heterodimer with ARNT and associated with other transcription factors such as COUP-TF, HIF, HNF4, LRH1, NRF1, PPAR, and RXR (Dere et al., 2011).

Analysis of published AhR ChIPseq data (Nault et al., 2016b) revealed HNF4α and COUP-TFII binding sited were over-represented within sequences under regions of enriched AhR, further alluding to interactions between the three transcription factors in regulating differential gene expression in the liver. Evaluation of differentially enriched binding sites suggests AhR co-operates with COUP-TFII and HNF4α enriched regions as suggested by the 47.85% and 70.09% of overlapping peaks with AhR, respectively. Due to ChIPseq limitations, it is not possible to definitively conclude if this involved physical interactions between AhR. COUP-TFII and/or HNF4α. Previous co-immunoprecipitation studies show COUP-TFI interacts with the AhR in MCF-7 cells and reported increased COUP-TFI expression inhibited TCDD-mediated luciferase activity under the control of the *CYP1A1* promoter (Klinge et al., 2000), an established AhR target gene. Furthermore, COUP-TFI binds to DREs (Klinge et al., 2000), the AhR binding motif.

Although previous results showed AhR and COUP-TFI (not COUP-TFII) cooperation, similar co-operative events are likely to occur between AhR and COUP-TFII. COUP-TFI and COUP-TFII share a high degree of protein homology with 98% similarity in the DNA binding domain and 96% in the ligand binding domain (Wang et al., 1991), suggesting they may possess overlapping protein interactions and functions (Polvani et al., 2019). For example, both COUP-TFI and COUP-TFII form heterodimers with the retinoid X receptor (RXR) to repress gene expression as well as compete for DNA binding with other transcription factors such as PPAR and HNF4 (Ashraf et al., 2019) (Kliewer et al., 1992; Kruse et al., 2008). Although COUP-TFs passively dysregulate HNF4α signaling events, direct interactions between COUP-TFII and HNF4α have been reported. COUP-TFII can either inhibit or promote HNF4α transactivation depending on the promoter, cell and tissue context (Schaeffer et al., 1993) (Stroup and Chiang, 2000). In the liver, COUP-TFII has been implicated in the regulation of lipid and glucose metabolism (Ashraf et al., 2019), while both HNF4α and COUP-TFII bind to the PPARα promoter region to impose opposite regulatory effects on β-oxidation (Pineda Torra et al., 2002). Specifically, HNF4α induces PPARα transcription, while COUP-TFII competes with HNF4α for PPAR regulatory region binding thereby causing PPARα repression (Pineda Torra et al., 2002). Consequently, dysregulation of HNF4α- and COUP-TFII-signaling is likely involved, albeit it through still to be determined mechanisms, in the progression of steatosis to hepatotoxicity and steatohepatitis with fibrosis by TCDD (Angrish et al., 2013; Nault et al., 2016c; Pierre et al., 2014).

HNF4α also serves a broader role beyond the regulation of hepatic metabolic events. The most prominent involve liver morphogenesis (Duncan et al., 1994; Parviz et al., 2003), and hepatocyte differentiation (Hayhurst et al., 2001; Morimoto et al., 2017). Previous studies suggest TCDD induces the loss of the liver-specific phenotype as evidenced by dysregulation of liver-specific gene expression (Nault et al., 2017a). In the present study, assessment of liver-specific DEGS following treatment with 30 μg/kg TCDD identified an association between reduced COUP-TFII binding and increased HNF4α binding in the presence of AhR genomic enrichment. Since HNF4α-signaling is required to maintain hepatic cell differentiation (Hayhurst et al., 2001), we evaluated the expression of genes previously identified to be dependent on this signaling pathway. These HNF4α-dependent genes in the liver were previously identified using a hepatocyte-specific HNF4α knockout model (Walesky et al., 2013). TCDD induced a cell de-differentiation phenotype, as evidenced by the overall repression of HNF4α-dependent gene expression involving a decrease in COUP-TFII binding with increased HNF4α binding. Loss of the hepatocyte phenotype was also evident in the overall repression of genes encoding the hepatic secretome. Similarly, repression of the hepatic secretome following TCDD treatment coincided with reduced COUP-TFII binding and increased HNF4α binding in the presence of AhR genomic enrichment. It is important to note, the hepatic secretome (Franko et al., 2019) only represents a set of secreted proteins from hepatocytes and does not account for secreted proteins produced by other hepatic cell types such as endothelial cells, hepatic stellate cells, cholangiocytes, and resident macrophages. Although loss of hepatocyte differentiation appears to occur, at least in part, indirectly through changes in HNF4α signaling, other mechanisms are also likely to contribute.

Hepatic cell de-differentiation following TCDD treatment plausibly occurs in part through changes in AhR signaling. Despite few studies linking AhR signaling to hepatic cell differentiation, AhR has been shown to regulate cell differentiation events in other tissues. For example, *in vitro* TCDD perturbed cardiomyocyte differentiation (Wang et al., 2013). Additionally, dysregulation of AhR-signaling has also been shown to promote proliferation and differentiation of human hematopoietic stem cells (HSCs) (Boitano et al., 2010). Under homeostatic conditions, HSCs remain in a quiescent state. However, following tissue damage or pathogen exposure HSCs are activated and subsequently proliferative. Continuously treated primary human CD34^+^ (a surface marker of HSCs) cells with StemRegenin 1, an AhR antagonist, for 53 weeks increased CD34^+^ cells by 73-fold (Boitano et al., 2010), further implicating AhR signaling in HSC differentiation. AhR deficient mice also exhibit elevated HSC proliferation with increased lymphocyte production and reduced levels of erythrocytes, neutrophils and monocytes (Singh et al., 2011). Interestingly, DREs are over-represented within 1,000 bps of HSC-enriched gene TSSs suggesting AhR regulation (Gazit et al., 2013). Moreover, AhR signaling plays a role stem cell differentiation into intestinal epithelial cells (Metidji et al., 2018). Although WNTβ-Catenin mediates intestinal stem cell proliferation and differentiation (Sato and Clevers, 2013), mice with dysregulated intestinal AhR signaling do not exhibit cell proliferation and intestinal crypt stem cell differentiation (Metidji et al., 2018). In the mouse liver, crosstalk between AhR and Wnt/β-catenin signaling has been reported (Schneider et al., 2014). TCDD also downregulates key Wnt/β-catenin pathway target genes in liver progenitor cells further implicating AhR signaling in hepatocyte differentiation. Collectively, these data suggest the AhR serves an important role in tissue-specific cell differentiation and is likely involved in hepatocyte differentiation.

In summary, metabolic reprogramming events caused by TCDD include dysregulation of the hepatic secretome as well as hepatocyte differentiation. These changes appear to involve intersections between AhR, HNF4α and COUP-TFII. Further studies are required to assess the co-operation between AhR, HNF4α and COUP-TFII, and to further define the types of interactions. Since the enrichment of AhR binding includes DNA motifs for a wide array of transcriptional regulators, it is also possible that TCDD-mediated hepatotoxicity occurs through mechanisms involving other transcription factors. The relevance of these interactions in human models also warrants further investigation.

## Notes

### Competing Interest Statement

The authors have declared no competing interest.

